# Tools and pipelines for BioNano data: molecule assembly pipeline and FASTA super scaffolding tool

**DOI:** 10.1101/020966

**Authors:** Jennifer M Shelton, Michelle C Coleman, Nic Herndon, Nanyan Lu, Ernest T Lam, Thomas Anantharaman, Palak Sheth, Susan J Brown

## Abstract

**Background:** Genome assembly remains an unsolved problem. Assembly projects face a range of hurdles that confound assembly. Thus a variety of tools and approaches are needed to improve draft genomes.

**Results:** We used a custom assembly workflow to optimize consensus genome map assembly, resulting in an assembly equal to the estimated length of the *Tribolium castaneum* genome and with an N50 of more than 1 Mb. We used this map for super scaffolding the *T. castaneum* sequence assembly, more than tripling its N50 with the program Stitch.

**Conclusions:** In this article we present software that leverages consensus genome maps assembled from extremely long single molecule maps to increase the contiguity of sequence assemblies. We report the results of applying these tools to validate and improve a 7x Sanger draft of the *T. castaneum* genome.

## Background

The quality and contiguity of genome assemblies, which impacts downstream analysis, varies greatly [1, 2, 3]. Initial assembly drafts, whether based on lower coverage Sanger or higher coverage next-generation sequencing (NGS) reads, are often highly fragmented. Physical maps of bacterial artificial chromosome (BAC) clones can be used to validate and scaffold sequence assemblies, but the molecular, human, and computational resources required to significantly improve a draft genome are often not available to researchers working on non-model organisms. The BioNano Irys ®System linearizes and images nicked and fluorescently labeled long DNA strands to generate single molecule physical maps. The Irys System provides affordable, high throughput physical maps of significantly higher contiguity with which to validate draft assemblies and extend scaffolds [4].

Genome assembly and scaffolding algorithms are inherently limited by the length of the DNA molecules used as starting material to generate data. Specifically, if repetitive, polymorphic or low complexity regions are longer than the single molecules used to generate data, then they cannot be resolved by bioinformatics tools with certainty. The specifications for PacBio P6-C4 chemistry [5] indicate that PacBio reads have an N50 of 14 kb with a maximum length of 40 kb. Illumina Long Distance Jump Libraries can also span 40 kb [6]. MinION nanopore sequence reads have an average read length of *<* 7 kb [7]. Illumina TruSeq synthetic long-reads can span up to 18.5 kb; however, they fail to assemble if the sequence has problematic regions longer than the component reads used to assemble the synthetic reads (e.g. in the heterochromatin) [8]. The OpGen Argus [9] platform produces optical maps that have a length of 150 kb to 2 Mb from up to 13 Gb data collected per run [10]. The Irys System from BioNano Genomics produces single molecule maps that have an average length of 225 kb from up to 96 Gb data collected per run after filtering for molecules *<* 150 kb [11]. Genomic repeats can be much longer than the 5-40 kb that many technologies can span with a single molecule. In fact, a recent study used consensus genome maps (assembled from single molecule maps *>* 150 kb) to identify repeats that are hundreds of kb in the human genome [12].

Sequence-based assembly methods are fraught with platform-specific error profiles (e.g. resolving homopolyer repeats or read-position effects on base quality) [13]. Map-based approaches offer an orthogonal genomic resource that complements sequence-based approaches but not their error profiles. For example, map-based error profiles tend to consider errors in estimated molecule or fragment length and errors associated with restriction sites that are too close together, neither of which influence sequence-based approaches [14, 12]. Both the BioNano Irys System and OpGen Argos platform provide single molecule maps from genomic DNA. OpGen may provide higher resolution maps by using enzymes with a six rather than a seven base pair recognition site, but BioNano’s single molecule maps still deliver a more efficient and affordable method for generating whole genome maps.

### Data formats

The tools described make use of three file formats developed by BioNano. The Irys System images extremely long molecules of genomic DNA that are nick-labeled at seven bp motifs using one or more nicking endonucleases and fluorescently labeled nucleotides. Molecules captured in TIFF images are converted to BNX format text files that describe the detected label position for each molecule (Figure 1(1-2)). The individual molecules described in BNX files are referred to as single molecule maps. Consensus Map (CMAP) files include the molecule map lengths and label positions for long genomic regions that are either inferred from assembly of raw single molecule maps (Figure 1(7-8)) or *in silico* from sequence scaffolds (Figure 1(3-4)). Individual maps in these two types of CMAP files are referred to as a consensus genome map or an *in silico* map, respectively. The alignment between two CMAP files is stored as an XMAP file that includes alignment coordinates and an alignment confidence score (Figure 1(10)).

**Figure 1.**
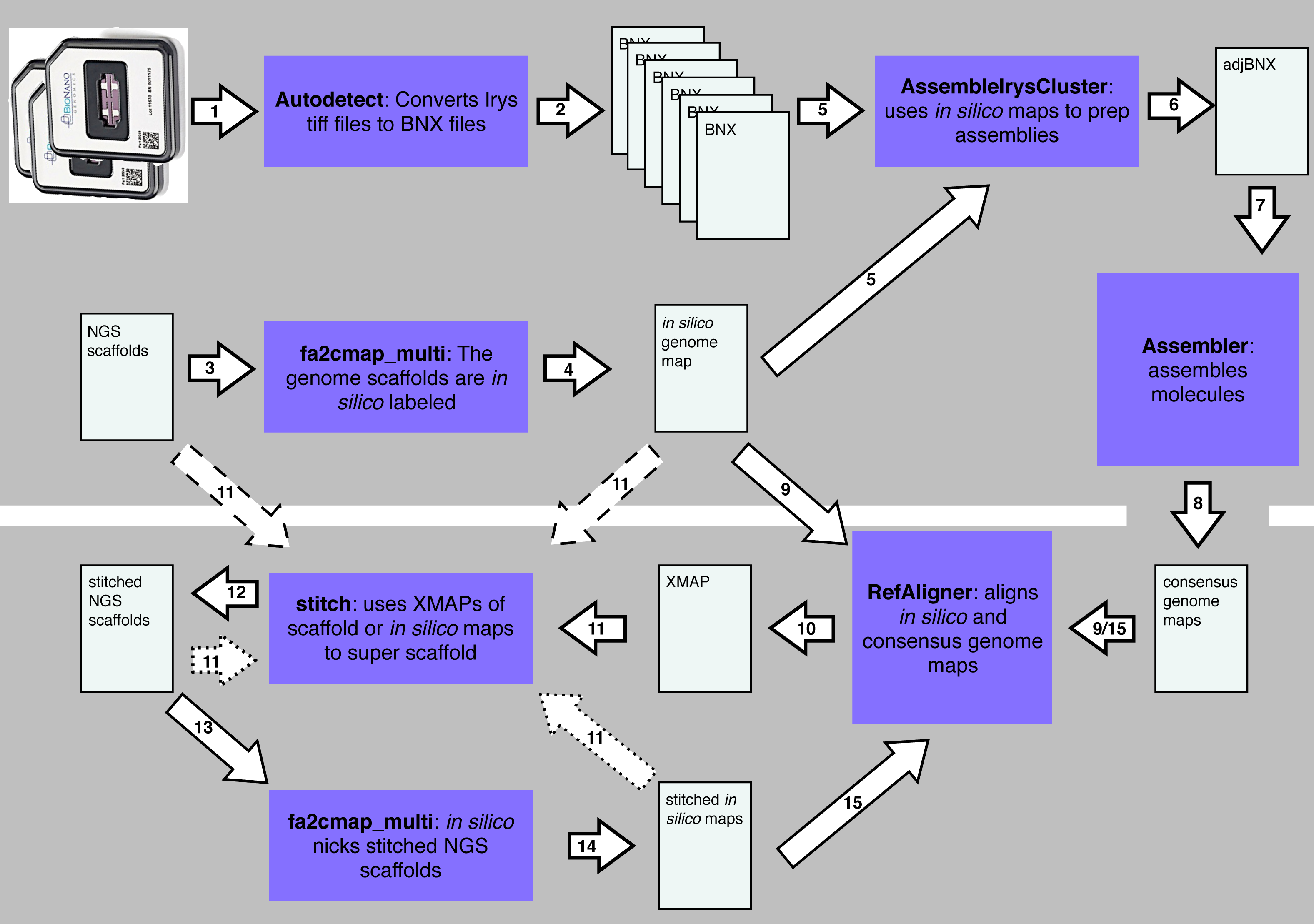
Data analysis steps. (1) Autodetect analyzes TIFF images of molecules and (2) outputs BNX text files. (3) Sequence scaffolds are digested *in silico* with fa2cmap multi producing (4) *in silico* maps. (5) AssembleIryscluster uses *in silico* maps, BNX files and estimated genome size to (6) normalize molecule stretch and set assembly parameters. (7) Assembler produces (8) a consensus genome map. (9) RefAligner aligns the consensus genome maps to the *in silico* maps producing (10) an XMAP alignment file. (11) XMAP, of *in silico* maps and consensus genome maps (see arrows with dashed lines) are used by stitch to produce (12) super scaffolded (stitched) sequence scaffolds. (13) Until no more super scaffolds are created the stitched sequence scaffolds are digested *in silico* with fa2cmap multi producing (14) a CMAP that is aligned to (9) the consensus genome maps and steps 10-15 are iterated. Arrows with dotted rather than dashed lines are used to as input during iterations.

### Other software tools for scaffolding with BioNano data

BioNano Genomics developed the Hybrid Scaffold tool to create more contiguous consensus genome maps using information from both sequence and BioNano genome map data. These more contiguous maps can then be used to create more contiguous sequence assemblies. The Hybrid Scaffold software first creates hybrid *in silico*/consensus genome map contigs based on an alignment between the two. The output genome maps are called hybrid scaffolds and are aligned to the original *in silico* maps. This alignment is used to output a FASTA file of sequence super scaffolds. These sequences include seven base pair ambiguous-base motifs to indicate where labels occur within gaps. Because they extend into regions with consensus genome maps but without sequence data they may begin or end with gaps. The Hybrid Scaffold program only generates hybrid *in silico*/consensus genome map contigs, and therefore super scaffolds, if no conflicts (e.g. negative gap lengths or otherwise conflicting alignments) are indicated in the alignment of *in silico* and consensus genome maps. In this conservative approach, all conflicting alignments are excluded from the hybrid scaffold genome map and flagged for further evaluation at the sequence level.

### Motivation

We designed tools and workflows to optimize the use of single molecule maps in the construction of whole genome maps and then use the best resultant consensus genome maps to improve contiguity of draft genome sequence assemblies. Single-molecule maps were assembled into BioNano genome maps *de novo* using software tools developed at BioNano [12]. As with sequence-based assembly algorithms, it was noted that testing a range of assembly parameters can improve final assembly quality for the BioNano Assembler. Additionally, applying error correction to molecule map stretch was found to improve assembly quality. Therefore, we created AssembleIrysCluster to normalize molecule map stretch and automate the writing of assembly scripts that use various parameters. We created the Stitch tool to super scaffold sequence-based assemblies using alignments to the optimal BioNano genome map. The Hybrid Scaffold and Stitch tools for genome finishing both take alignments from the BioNano RefAligner as input. Both tools were developed simultaneously but were ultimately found to be useful for distinct applications. We validated AssembleIrysCluster and Stitch using the *Tribolium castaneum* genome [15] because this project has genetic map resources [16] that offer independent corroboration. Genetic maps were not used as input for Stitch. In this case, the super scaffolds created by Stitch were compared to the order of scaffolds within ChLGs predicted by the genetic map.

## Implementation

### Overview

The tools described below take raw molecule maps as input, assemble genome maps, and then use these genome maps to super scaffold draft sequence assemblies. The tool AssembleIrysCluster generates consensus genome maps for a range of assembly parameters. We developed AssembleIrysCluster to prepare BNX files for assembly and produce nine customized assembly scripts (Figure 2C-G). Next, genome maps from the user-selected best assembly are used by the tool Stitch to validate and super scaffold sequence assemblies (Figure 3).

**Figure 2.**
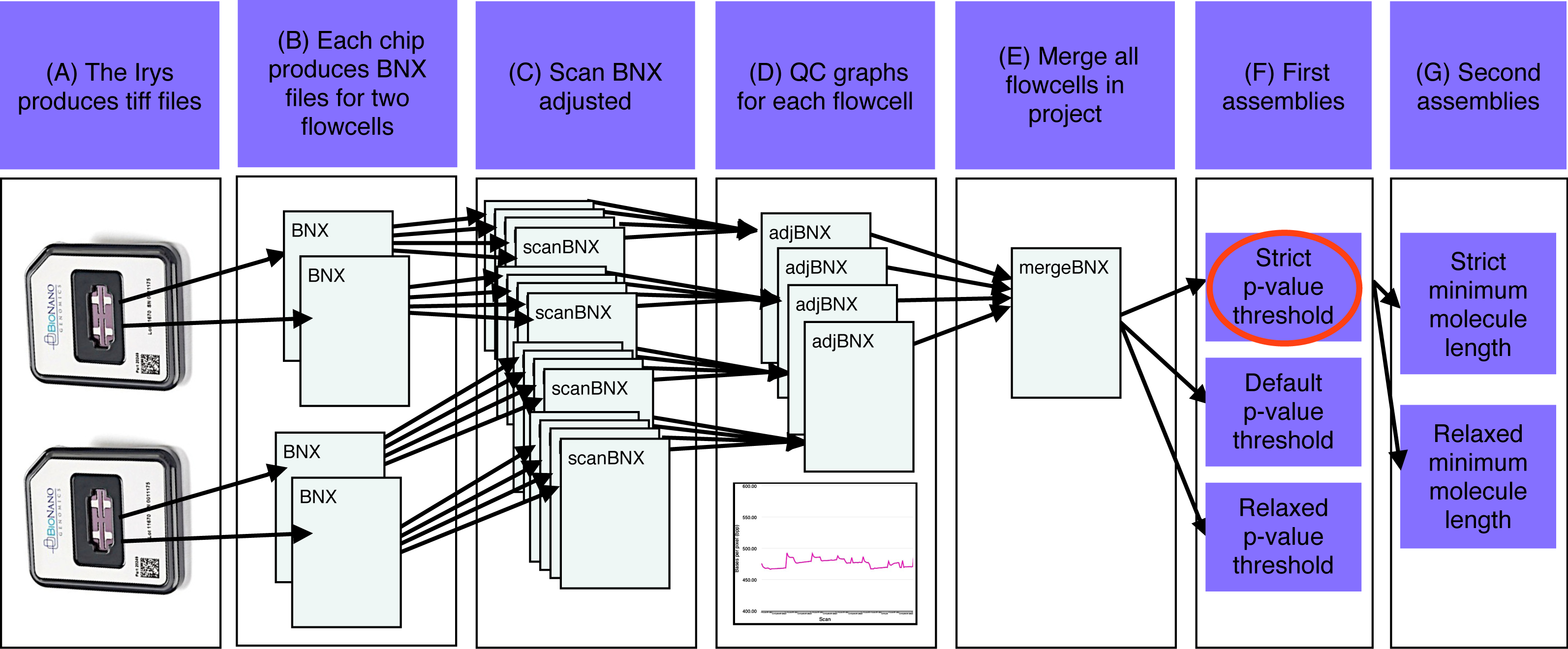
Assembly workflow for assemble SGE cluster.pl. (A) The Irys instrument produces tiff files that are converted into BNX text files. (B) One BNX file is produced for each flowcell on a IrysChip. (C) BNX files are split by scan and aligned to the sequence reference. Stretch (bases per pixel) is recalculated for each scan from the alignment. (D) Quality check graphs are created for each pre-adjusted flowcell BNX. (E) Adjusted flowcell BNXs are merged. (F) The first assemblies are run with a variety of p-value thresholds. (G) The best first assembly (red oval) is chosen and a version of this assembly is produced with a variety of minimum molecule length filters.

**Figure 3.**
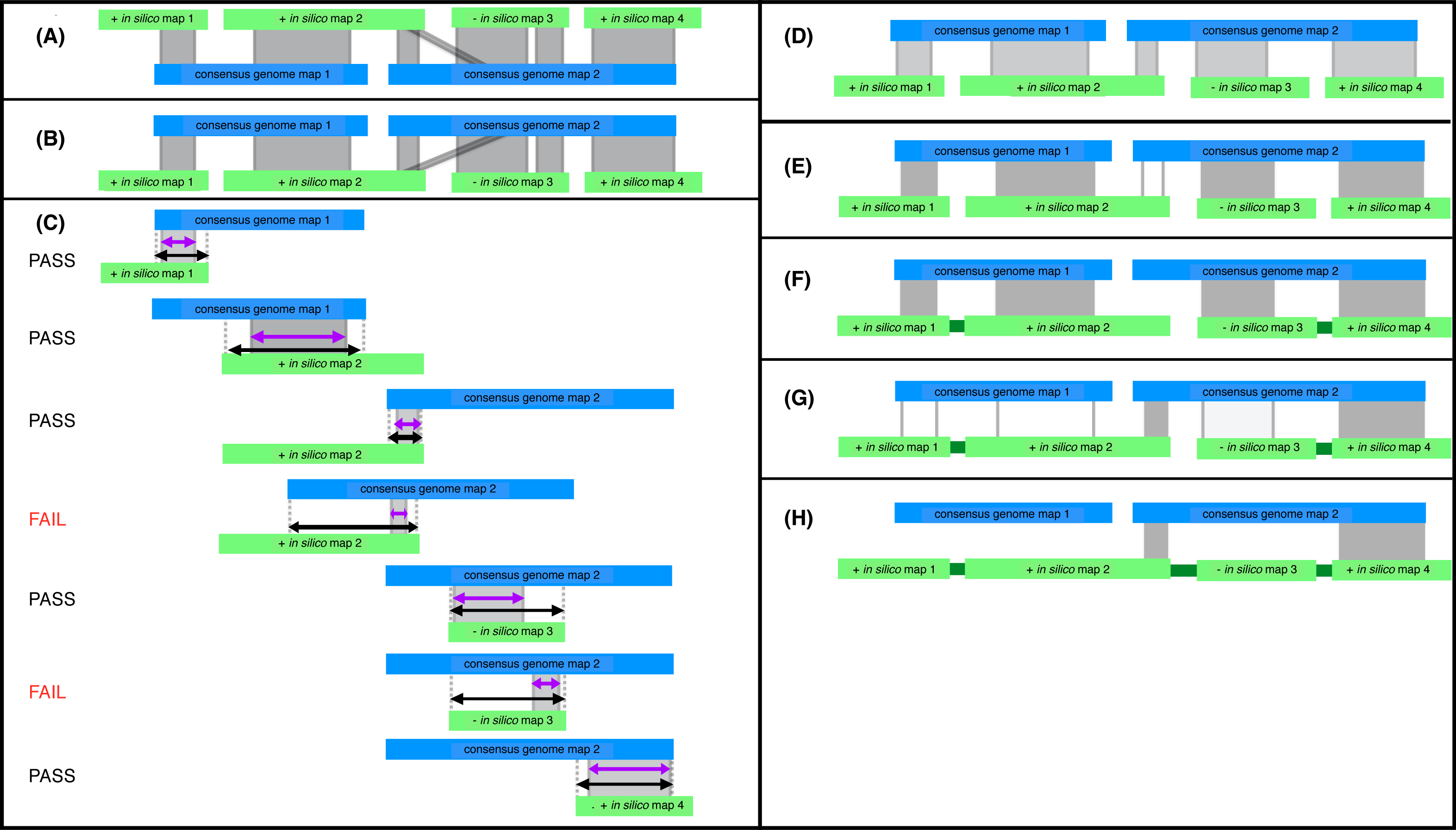
Steps of the stitch.pl algorithm. Consensus genome maps (blue) are shown aligned to *in silico* maps (green). Alignments are indicated with grey lines. CMAP orientation for *in silico* maps is indicated with a ”+” or ”-” for positive or negative orientation respectively. (A) The *in silico* maps are used as the reference. (B) The alignment is inverted and used as input for stitch.pl. (C) The alignments are filtered based on alignment length (purple) relative to total possible alignment length (black) and confidence. Here assuming all alignments have high confidence scores and the minimum percent aligned is 30% two alignments fail for aligning over less than 30% of the potential alignment length for that alignment. (D) Filtering produces an XMAP of high quality alignments with short (local) alignments removed. (E) High quality scaffolding alignments are filtered for longest and highest confidence alignment for each *in silico* map. The third alignment (unshaded) is filtered because the second alignment is the longest alignment for *in silico* map 2. (F) Passing alignments are used to super scaffold (captured gaps indicated in dark green). (G) Stitch is iterated and additional super scaffolding alignments are found using second best scaffolding alignments. (H) Iteration takes advantage of cases where *in silico* maps scaffold consensus genome maps as *in silico* map 2 does. Stitch is run iteratively until all super scaffolding alignments are found.

### AssembleIrysCluster: Molecule Stretch

In the first stage, AssembleIrysCluster adjusts single molecule map stretch (Figure 2C). BioNano software operates under the assumption that imaged molecules contain 500 bases per pixel (bpp). Stretch, or bpp, can deviate from 500 bpp and this discrepancy can vary from scan to scan within a flowcell (Additional file 1). Sequence scaffolds are considered to be more accurate than raw BNX molecules in terms of label positions. Therefore molecule maps in BNX files are split by scan, and after alignment to the *in silico* maps, an empirical average bpp value is determined for the molecule maps in each scan. The bpp indicated by this alignment is used by RefAligner to adjust molecule map bpp to 500. Once stretch has been evaluated and normalized, the split BNX files are merged into a single file (Figure 2E).

We observed consistent patterns of empirically determined bpp between flowcells using the same flowcell model and chemistry when signal-to-noise ratios are optimal and the degree of genomic divergence between the samples used for the *in silico* maps and molecule maps are low (Additional file 1). To identify the low quality flowcells, bpp observed in alignments of scans are plotted as a quality control (QC) graph by AssembleIrysCluster (Figure 2D).

### AssembleIrysCluster: Customization of BioNano Assembly Scripts

In the next stage, AssembleIrysCluster creates various assembly scripts to explore a range of parameter sets around the the assembler default parameters with the goal of selecting the optimal assembly for downstream analysis (Figure 2F-G). Single molecule maps in the adjusted merged BNX file are aligned to the *in silico* maps. An alignment error profile generated by RefAligner is used with the estimated genome length to calculate default assembly parameters; and the eight other scripts that include variants of these parameters. Initially, three assemblies are run, the first with 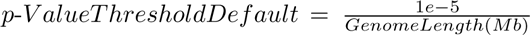 the second with 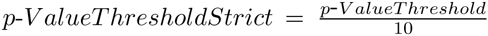 and the third with *p*-*ValueThresholdRelaxed* = *p*-*ValueT hreshold* × 10 (Figure 2F). The minimum molecule length is set to 150 kb. If one of these runs does not produce a satisfactory assembly, then two minimum molecule length variants (180 kb and 100 kb) are tested with the *p*-*ValueThreshold* of the current best assembly (Figure 2G). Between three and nine assemblies are run until a satisfactory assembly is produced.

### BioNano Assembly Optimization

The ultimate goal is to produce consensus genome maps that can be used to guide sequence-based haploid reference genome assembly. While single molecule maps can be used to reconstruct haplotypes [17], genome assembly involves collapsing polymorphisms arbitrarily into a consensus reference genome. Therefore the cumulative length of ideal consensus genome maps should equal the haploid genome length. Additionally, 100% of the consensus genome maps would align non-redundantly to 100% of the *in silico* maps. In practice, the best BioNano assembly is selected based on similarity to the estimated haploid genome length and minimal alignment redundancy to the reference *in silico* maps.

### Stitch: Alignment Filters

For each RefAligner alignment the user designates a reference file with *in silico* maps from either a draft or reference quality genome or, if the user is aligning two files that both contain consensus genome maps, then a file of consensus genome maps. Additionally, the user specifies a query file with either single molecule maps or consensus genome maps.

We designed the Stitch algorithm to use alignments of query consensus genome maps to reference *in silico* maps in order to predict the higher order arrangement of sequence scaffolds (Figure 3). RefAligner assumes the alignment reference has the error profile of *in silico* maps and the query has the error profile of consensus genome maps. Therefore alignments are run with the *in silico* maps as the reference, and are inverted and sorted by consensus genome map coordinates for efficient parsing by Stitch (Figure 3A-B).

Before inferring super scaffolds from XMAP files, Stitch filters low quality alignments by confidence score. Alignments of *in silico* and consensus genome maps are assigned a confidence score that is the −*log*10 of the *False Positive p*-*Value*. Misaligned labels and sizing error increase the alignment *False Positive p*-*Value* and decrease confidence scores [18].

Super scaffolds are built from overlapping alignments. Overlapping alignments are similar to global alignments, i.e., alignments spanning from end to end for two maps of roughly equal length, but to search for overlap alignment gaps after the ends of either map are not penalized. The RefAligner scoring scheme does not currently have a parameter to favor overlapping alignments, e.g., to initialize the dynamic programming matrix with no penalties and take the maximum score of the final row or column in the matrix. RefAligner reports local alignments between two maps and applies a fixed penalty based on the user-defined likelihood of unaligned labels at the ends of the alignment. Raising or lowering this penalty selects for local or global alignments, respectively, but neither option favors overlapping alignments specifically. Stitch filters by the percent of the total possible alignment length that is aligned (Figure 3C). To approximate scoring that favors overlapping alignments, Stitch uses thresholds for minimum percent of total possible aligned length, the percent aligned threshold (PAT).

Similar to scoring structures that favor overlapping alignments, PAT filters out local alignments. However, unlike a scoring structure, PAT is applied after alignment and therefore cannot result in the aligner exploring possible extensions into an overlap but instead favors a shorter local alignment with a higher cumulative score. Therefore Stitch accepts alignments with less than 100% PAT. Default values for the PAT were determined empirically after reviewing the degree to which filtered alignments agreed with the independently derived genetic maps of *T. castaneum* and by visual inspection of alignments.

In practice we used two sets of alignment filters and kept alignments that passed one or both sets. The first set had a low PAT and a high confidence score threshold. The second set had a higher PAT and a lower confidence score and was intended to identify longer overlaps especially in regions of the genome where label density is low.

### Stitch: Super scaffolding

Scaffolding alignments are selected from the remaining high quality alignments (i.e., more than one *in silico* map aligns to the same consensus genome map (Figure 3D)). For each *in silico* map with more than one high quality, scaffolding alignment the longest alignment for the *in silico* map is selected (Figure 3D-E). If alignment length is identical then the highest confidence alignment is selected. If confidence scores are identical then an alignment is chosen arbitrarily.

Gap lengths between *in silico* maps are inferred from scaffolding alignments and used to create new super scaffolds (Figure 3F) in a new genome FASTA file and associated AGP file. If gap lengths are estimated to be negative, Stitch adds a 100 bp spacer gap to the sequence file and indicates that the gap is type ”U” for unknown in the AGP.

Stitch only makes use of one alignment per *in silico* map each iteration. Stitch can be run iteratively (Figure 1(10-15)) such that each successive output FASTA file is converted into *in silico* maps and aligned to the original consensus genome maps. This alignment is inverted and used as input for the next iteration. Subsequent iterations of Stitch will make use of any *in silico* maps that join growing super scaffolds, effectively using both sequence data and genome maps to stitch together the final super scaffolds (Figure 3G-H).

### Stitch: Flagging Potential Mis-assemblies

This algorithm is meant to be an intermediate refinement of draft genomes prior to further fine scale refinement at the sequence level. Inconsistencies between the consensus genome maps and the *in silico* maps are reported in output logs to facilitate downstream sequence editing. If an alignment passes initial confidence score and PAT filtering but has a PAT less than 60%, this is reported as a partial alignment. A partial alignment may occur if either the sequence scaffold or the consensus genome map is the result of a chimeric or erroneous assembly. Additionally, if a gap length is estimated to be negative, it may indicate that the sequence scaffolds can be joined with a local assembly or that an incorrect assembly needs to be broken within a scaffold. Assembly errors in the consensus genome maps or spurious alignments could also result in either of these cases. Ideally researchers could make use of the alignment of genomic sequence reads to the genome sequence assembly and the alignment of single molecule maps to the consensus genome maps to determine which assembly is likely to be incorrect.

### Post Analysis: Software Updates

Since completion of the *T. castaneum* genome update we have released updates for both Stitch and AssembleIrysCluster. Stitch (version 1.4.5) now allows the user to set a minimum negative gap length filter for alignments. In the event that two *in silico* maps have a negative gap length smaller than this value, which is equivalent to a longer overlap of the sequence scaffolds, Stitch will automatically exclude both *in silico* maps from consideration when super scaffolding. This new feature allows users to further customize the stringency of the Stitch output. In addition, BioNano Genomics has updated Assembler to automate per-scan stretch adjustment. Finally the KSU K-INBRE Bioinformatics Core has moved to producing assemblies on a Xeon Phi server with 1488 threads (6x60x4 Xeon Phi co-processor threads + 24×2 Xeon host threads) and 256GB of host RAM + 6 x 8GB Xeon Phi Ram, and Linux CentOS 7 operating system. Because of all of these changes we opted to create a new assembly workflow rather than update AssembleIrysCluster. The most current assembly workflow, AssembleIrysXeonPhi maintains all the functionality of AssembleIrysCluster (e.g. adjusting stretch by scan and writing assembly scripts with all combinations of three *p*-*ValueThresholds* and three *Minlen* parameters) but runs on our new machine with the latest release of the BioNano Assembler and RefAligner.

### Post Analysis: Tutorials and Complete Pipelines

Additionally, since completion of the *T. castaneum* genome update we have created three pipelines for BioNano data complete with sample datasets and tutorials. The Sewing machine pipeline iteratively super scaffolds genome FASTA files with Bio-Nano genome maps using stitch.pl and the BioNano tool RefAligner until no new super scaffolds can be produced. The pipeline runs alignments with both default and relaxed parameters. These alignments are then used by stitch.pl to superscaffold a fragmented genome FASTA. The ”Raw data-to-finished assembly and assembly analysis” pipeline for BioNano molecule maps with a sequence-based genome FASTA prepares raw molecule maps and writes and runs a series of assemblies for them. Then the user selects the best assembly and uses this to super scaffold the reference FASTA genome file and summarize the final assembly metrics and alignments. The ”Raw data-to-finished de novo assembly and assembly analysis” pipeline for BioNano molecule maps is for *de novo* projects. Both of the last two pipelines are broken into stages at points were the user must select the best set of parameters from assemblies or alignments that have been run by the pipeline.

## Results and Discussion

### Dataset generation

High molecular weight DNA was isolated from young *Tribolium castaneum* pupae from the GA2 line that was inbred 20 generations. The GA2 line was also used for the genome sequence-based assembly. Ethical approval was not sought for the study because the study organism is an insect.

Using Knickers (BioNano Genomics), an *in silico* label density calculator, we estimated that the *Tribolium castaneum* genome had 8.66 and 5.51 labels per 100 kb for the nt.BspQI and nt.BbvCI enzymes (New England BioLabs) respectively. The ideal number of labels per 100 kb is between 10 and 15 therefore we nicked with both enzymes. DNA was nicked, labeled with fluorescent nucleotides, and repaired according to BioNano protocol; and 93 BNX files were produced from the Irys genome mapping system (Figure 4 and Additional file 2). Four corrupted files (cumulative length = 0) were excluded from this analysis. All *T. castaneum* BNX files have been deposited to labarchives (doi:10.6070/H4V69GK3).

**Figure 4.**
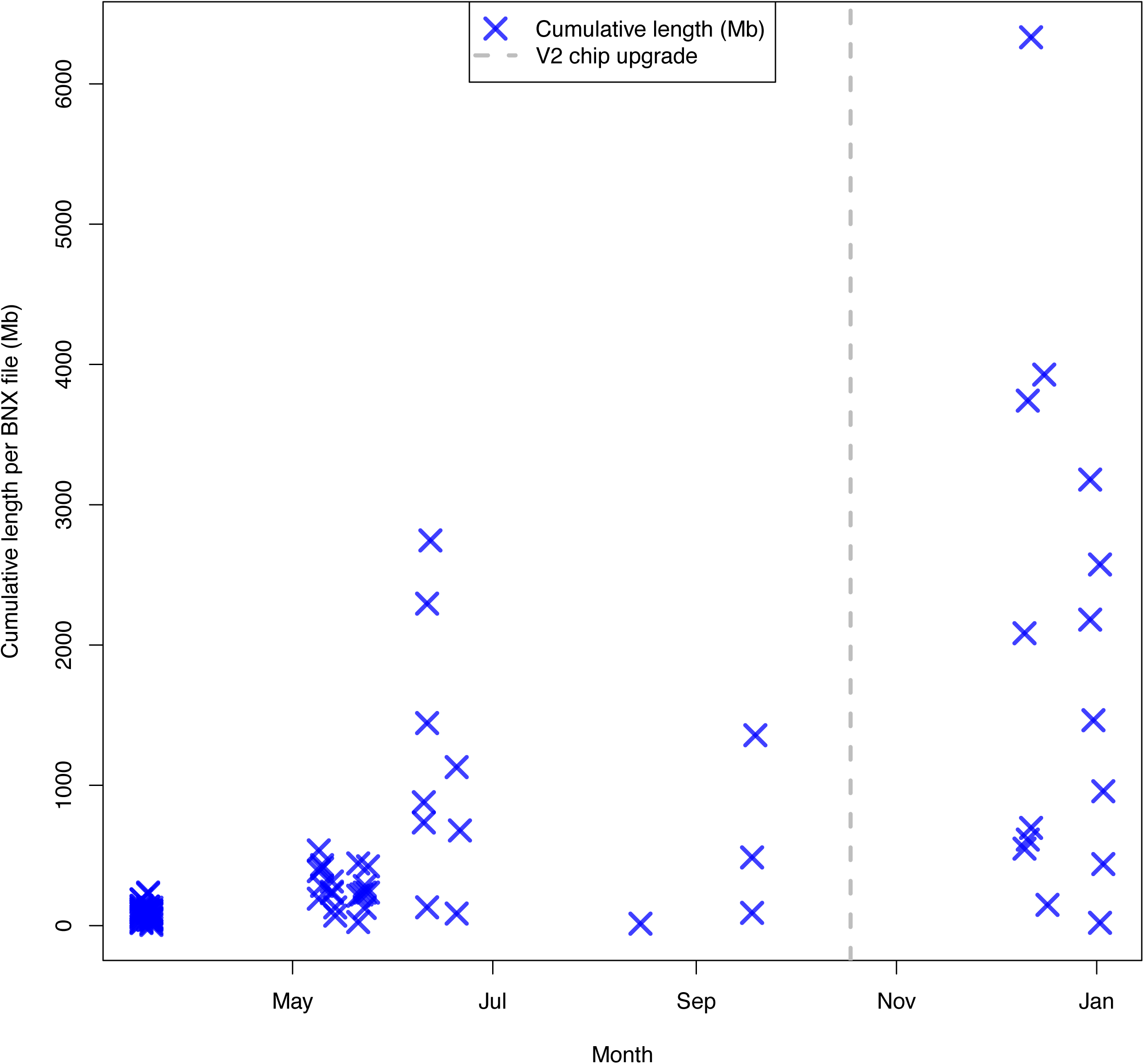
Cumulative length per BNX file for T. castaneum data generated over time. Cumulative length of single molecule maps > 150 kb are plotted on the y-axis (purple X), the upgrade to the V2 IrysChip (grey dashed line) is plotted and date is indicated on the x-axis. Data was generated from 03-2013 to 01-2014. Aborted runs (cumulative length = 0) excluded.

The high number of BNX files produced is due to several factors. Typically one BNX file is produced per flowcell; however, in certain cases after the initial number of scans, additional scans were run producing an additional BNX file. Another reason for the large number of files is that data was originally generated using the IrysChip® V1 while the Irys System was under beta testing. Over time, maximum cumulative length per BNX file increased (Figure 4 and Additional file 2). After the upgrade to the IrysChip V2, an increase was also observed in maximum data per BNX file (Figure 4 and Additional file 2).

Molecule map quality metrics were calculated using bnx stats (version 1.0). We generated ∼250x coverage of the *T. castaneum* genome for single molecule maps *>*150 kb, the default minimum molecule map length. The 239,558 single molecule maps with lengths *>*150kb had an N50 of 202.63 kb and a cumulative length of 50.6 Gb (Table 1). Histograms of per-molecule quality metrics for maps after applying a minimum length filter of either 100 kb, 150 kb and 180kb are reported in Additional file 3.

**Table 1.** Single molecule maps from *T. castaneum* filtered by minimum length. Molecule map N50, cumulative length and number of maps are listed for all three molecule length filters for the *T. castaneum* genome data.

| Minimum molecule map length (kb) | Molecule map N50 (kb) | Cumulative length (Mb) | Number of molecule maps |
| --- | --- | --- | --- |
| 100 | 165.35 | 82,738.71 | 503,414 |
| 150 | 202.64 | 50,579.12 | 239,558 |
| 180 | 232.57 | 34287.15 | 139,949 |

To generate the *in silico* maps, we used the sequence-based assembly scaffolds from version 5.0 of the *Tribolium castaneum* genome (Tcas5.0). Version 5.0 (Table 2) of the *T. castaneum* genome is an updated version of the sequence assembly that was created by adding 1.03 Mb of sequence to version 3.0 [15]. Two hundred and twenty-three of the 2240 scaffolds within Tcas5.0 were longer than 20 kb with more than 5 labels, the minimum requirements for an *in silico* map. These longer sequence scaffolds represent the bulk of the sequence assembly, 152.53 of the 160.74 Mb (Table 2).

**Table 2.** *T. castaneum* assembly summary. Assembly metrics for Tcas5.0 (the starting sequence scaffolds), the Tcas5.0 *in silico* maps, the consensus genome map of assembled molecule maps, the automated output of Stitch (Tcas5.1), the manually curated sequence assembly (Tcas5.2) and the sequence assembly produced by the BioNano Hybrid Scaffold software for the *T. castaneum* genome.

|  | N50 (Mb) | Number | Cumulative Length (Mb) |
| --- | --- | --- | --- |
| Tcas5.0 sequence scaffolds | 1.16 | 2240 | 160.74 |
| Tcas5.0 *in silico* maps | 1.20 | 223 | 152.53 |
| Consensus genome maps | 1.35 | 216 | 200.47 |
| Tcas5.1 sequence scaffolds | 3.85 | 2148 | 165.72 |
| Tcas5.2 sequence scaffolds | 4.46 | 2150 | 165.92 |
| Tcas BioNano hybrid scaffolds | 1.83 | 2210 | 175.54 |

### Assembly: Selecting the Optimal BioNano Assembly

Single molecule maps were assembled *de novo* into five distinct BioNano genome maps for *T. castaneum*.

Molecule maps were prepared for assembly (Figure 2C) and noise parameters were estimated (Figure 2D-E) using AssembleIrysCluster (version 1.0). Specifically, AssembleIrysCluster calculated three ”-T” parameters (5e-10, 5e-09 and 5e-08) based on the estimated genome size, and adjusted molecule map stretch and estimated molecule map noise parameters based on alignment to *in silico* maps. Assemblies with these p-value thresholds are named *Relaxed*-*T*, *Default*-*T* and *Strict*-*T*, respectively.

In the first round of selection, the *Strict*-*T* assembly was the best of these three assemblies because it has a cumulative size close to 200 Mb (Table 2 and Figure 5), the estimated size of the *T. castaneum* genome, and a small difference between non-redundant aligned length or breadth of alignment, and total aligned length (Figure 5, Table 3 and Additional file 4). Thus in the second round of selection, *Strict*-*T* parameter was used for two further assemblies that had relaxed minimum molecule length (*Relaxed*-*Minlen*) of 100 kb rather than the 150 kb default or a strict minimum molecule length (*Strict*-*Minlen*) of 180 kb. The *Strict*-*T AndStrict*-*Minlen* assembly improved alignments by reducing redundancy slightly. However the cumulative length of the assembly was 21.45 Mb smaller than the estimated genome size. The *Strict*-*T AndRelaxed*-*M inlen* assembly had a worse cumulative length and alignment redundancy than the *Strict*-*T* assembly. Because neither of the assemblies using 100 or 180 kb as the minimum molecule length improved both assembly metrics when compared to the *Strict*-*T* assembly, generated in the first round of selection with the default *Minlen* of 150 kb, the *Strict*-*T* assembly will be referred to as the *T. castaneum* consensus genome maps in further analysis.

**Table 3.**
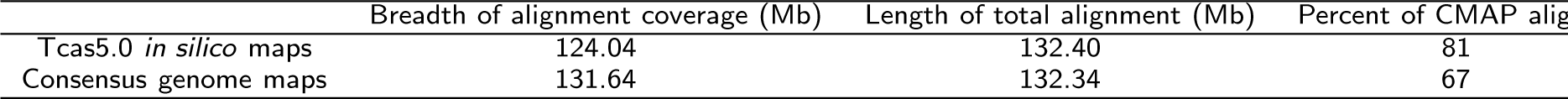
Alignment of *T. castaneum* consensus genome maps to the *in silico* maps of Tcas5.0. Breadth of alignment coverage (non-redundant alignment), length of total alignment (including redundant alignments) and percent of CMAP covered (non-redundantly) were calculated for the *in silico* maps and the consensus genome maps of the *T. castaneum* genome the using xmap stats.pl.

|  | Breadth of alignment coverage (Mb) | Length of total alignment (Mb) | Percent of CMAP aligned |
| --- | --- | --- | --- |
| Tcas5.0 *in silico* maps | 124.04 | 132.40 | 81 |
| Consensus genome maps | 131.64 | 132.34 | 67 |

**Figure 5.**
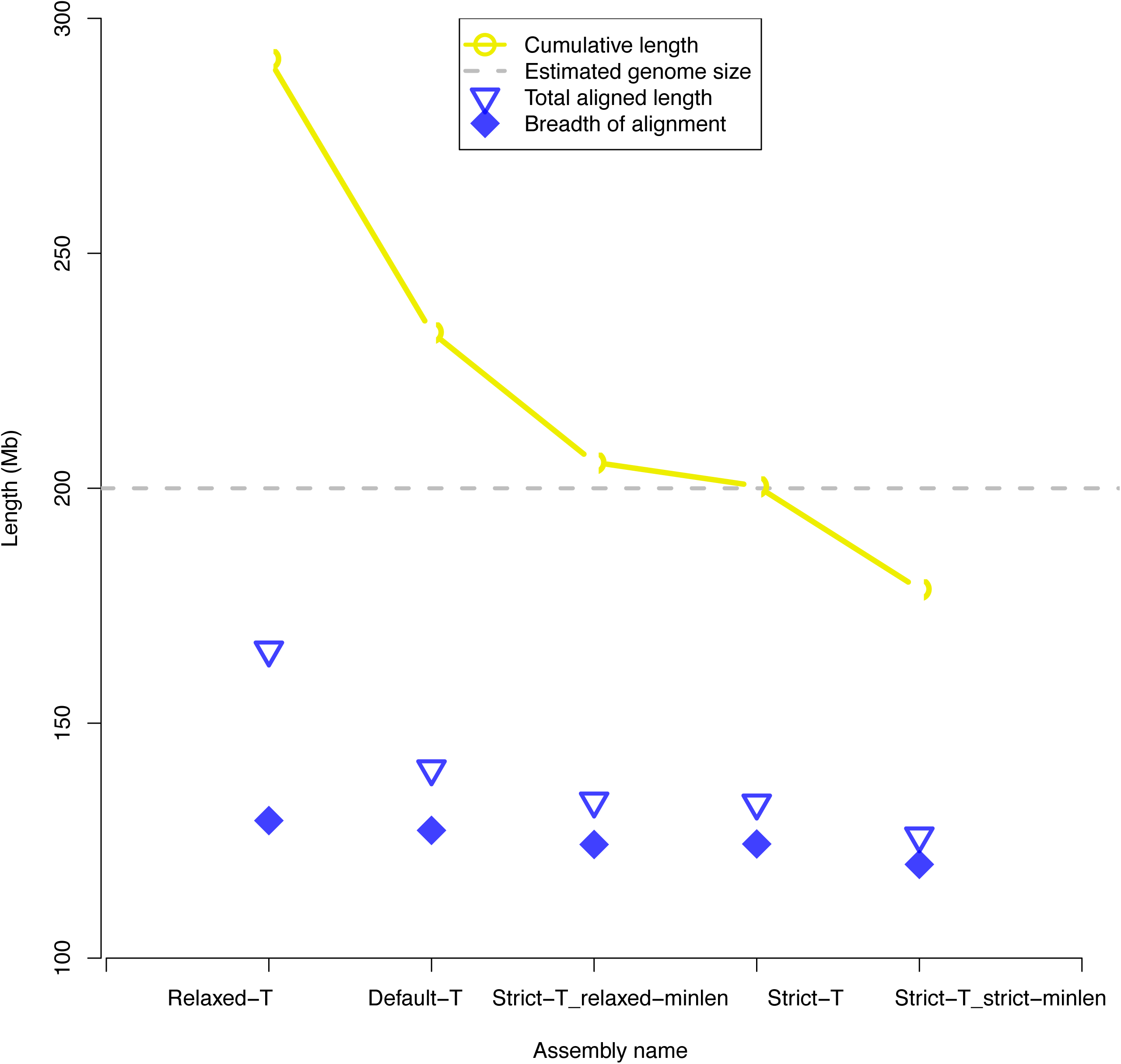
Comparison of assemblies from the T. castaneum single molecule maps using five sets of parameters. Relaxed, default and strict ”-T” parameters were set to 5e-08, 5e-09 and 5e-10. Relaxed, default and strict minimum molecule length were set to 100, 150 and 180 kb.

The *T. castaneum* consensus genome maps have an N50 of 1.35 Mb, a cumulative length of 200.47 Mb, and 216 genome maps (Table 2 and Figure 5). *T. castaneum* consensus genome maps aligned to 124.04 Mb of the *in silico* maps created from the Tcas5.0 assembly validating 81% of the draft sequence assembly (Table 3 and Figure 5). Assembly metrics were calculated using the BNGCompare script (version 1.0). More detailed assembly metrics for all five assembled consensus genome maps are available in Additional file 4. Assembled *T. castaneum* genome maps have been deposited to labarchives (doi: 10.6070/h42f7kf2).

### Stitch: Automated and Manually Edited Assemblies

Tcas5.1 is the output of Stitch (version 1.4.4) run for five iterations with two sets of alignment filters. To select quality alignments from regions of high and low label density, the first minimum confidence was 13 and the *PAT* was 30 and the second minimum confidence was 8 and the *PAT* was 90. The resulting super scaffolds showed a greater than three-fold increase in N50 from 1.16 to 3.85 Mb (Table 2).

The Tcas5.1 super scaffolds captured an additional 92 gaps between Tcas5.0 sequence scaffolds. Sixty-six gaps were estimated to have positive gap lengths and were represented in the sequence assembly with their estimated size (Figure 6). Twenty-six gaps were estimated to have negative lengths and were represented with spacers of 100 N’s (Figure 6). Extremely small negative gap lengths (*<* −20 kb) were flagged for further evaluation at a sequence level. In some cases, extremely small negative gaps lengths suggest that a chimeric sequence scaffold may need to be broken at the sequence level and its fragments incorporated into different chromosome linkage groups (ChLGs). For example, half of scaffold 81 from Tcas5.0 aligns between scaffolds 80 and 82 on ChLG5 (Figure 7A) while the other half aligns between scaffolds 102 and 103 from ChLG7 (Figure 7B). Scaffold 81 from Tcas5.0 was placed in ChLG5 in the *T. castaneum* genetic map. The arrangement supported by the genetic map was selected for cases like this where manual editing was required. The manually curated assembly is referred to as Tcas5.2. In Tcas5.2, joins were also manually accepted if they agreed with the genetic map but the alignment quality was low. Ultimately, Tcas5.2 had a higher N50 than the automated Stitch output, 4.46 Mb (Table 2), with 66 gaps with positive estimated lengths and 24 negative length gaps.

**Figure 6.**
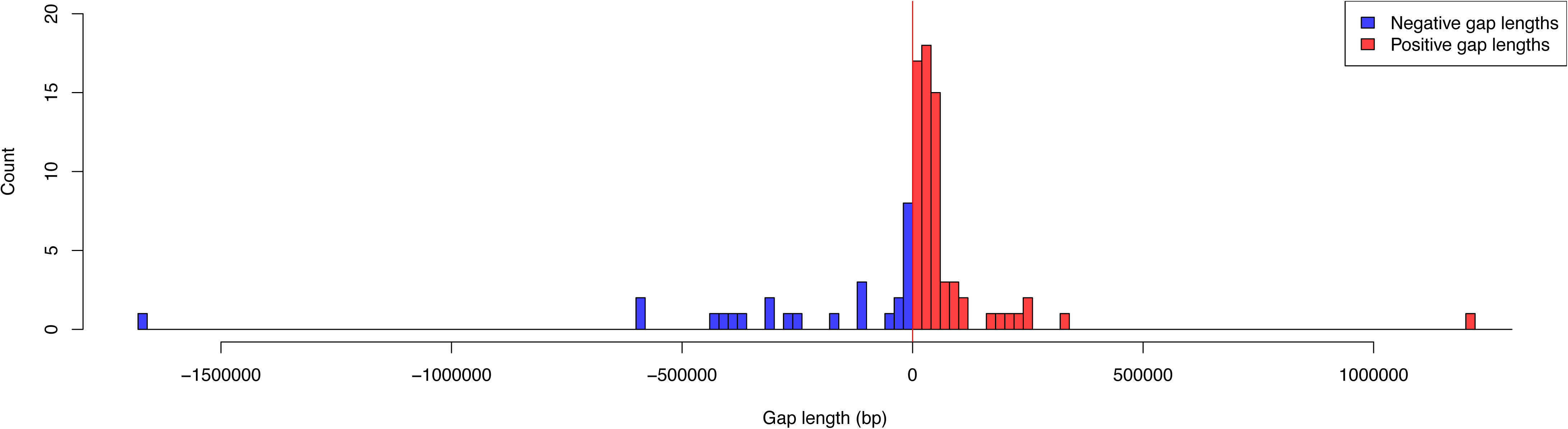
Histogram of gap lengths in Tcas5.1. Positive and negative gaps lengths for Tcas5.1 added to the automated output of stitch.pl based on filtered scaffolding alignments. The majority of gap lengths added by stitch.pl, 66, were positive (red). The remaining 26 gaps had negative lengths (purple).

**Figure 7.**
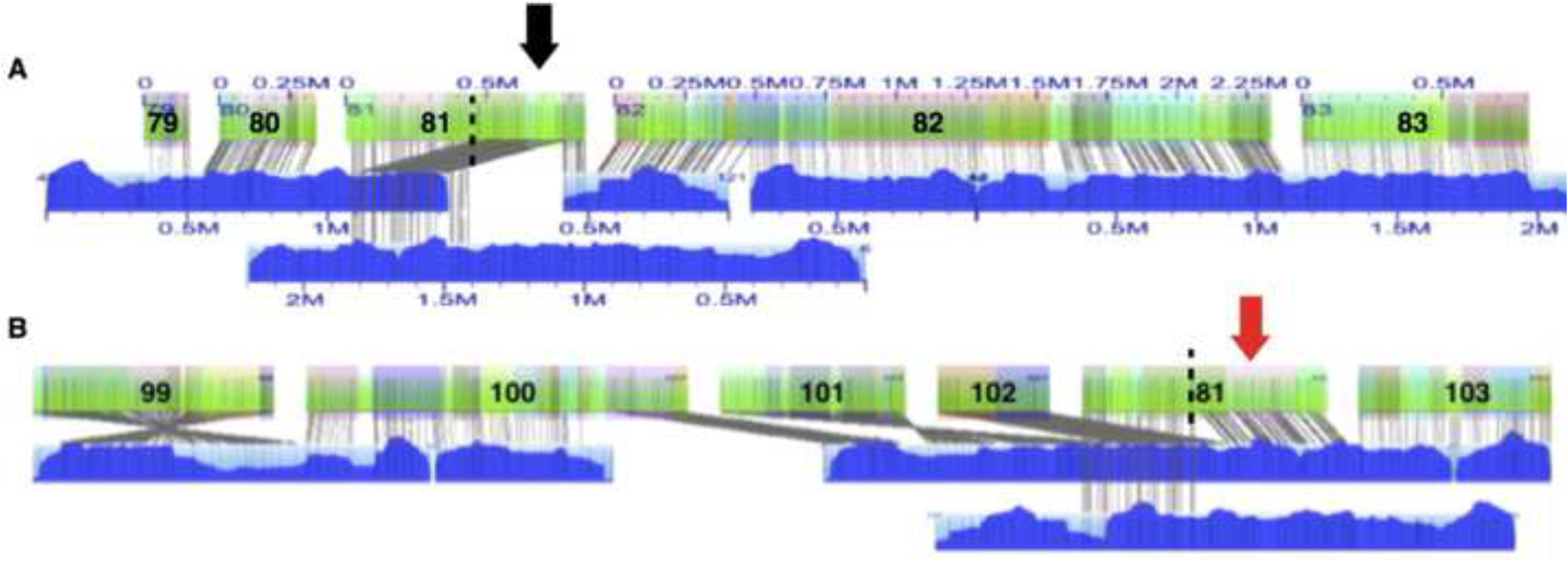
Extremely small negative gap length for *in silico* map of scaffold 81. Two XMAP alignments for *in silico* map of sequence scaffold 81 are shown. Consensus genome maps (blue with molecule map coverage shown in dark blue) align to the *in silico* maps of scaffolds (green with contigs overlaid as translucent colored squares). Sequence scaffolds 79-83 were placed within ChLG 5 and sequence scaffolds 99-103 were placed with ChLG 7 by the *T. castaneum* genetic map. (A) Half of the *in silico* map of sequence scaffold 81 aligns with its assigned ChLG (black arrow). (B) The other half aligns with ChLG 7 (red arrow) producing a negative gap length smaller than -20 kb. The alignment that places sequence scaffold 81 with ChLG 7 disagrees with the genetic map and was manually rejected for Tcas5.2.

Nearly every ChLG was less fragmented in the Tcas5.2 assembly than in the Tcas5.0 assembly. The number of scaffolds was reduced for each ChLG (Table 4) except for the 26 relatively short (N50 = 0.05 Mb) and unlocalized scaffolds from ChLGY. For example, ChLGX was reduced from 13 scaffolds (Figure 8B) to 2 in the final super scaffold or chromosome build (Figure 8A). Five scaffolds were reoriented in ChLGX based on alignment to the consensus genome maps (Figure 8A-B). Scaffolds were also reordered based on alignment to consensus genome maps. For example, scaffold 2 from ChLGX aligned between scaffold 36 and 37 of ChLG 3 and was therefore moved in Tcas5.2 (Additional file 5). Also, 19 previously unplaced scaffolds were anchored within a ChLG (Table 4). Improvements in Tcas5.2 over Tcas5.0 are shown in alignments of the respective *in silico* maps to consensus genome maps for all ChLGs in Additional file 5.

**Table 4.** Each *T. castaneum* chromosome linkage group (ChLG) before and after super scaffolding. The number of sequence scaffolds in the ordered Tcas5.0 ChLG bins and the number of sequence super scaffolds and scaffolds in the Tcas5.2 ChLG bins. The number of sequence scaffolds that were unplaced in Tcas5.0 and placed with a ChLG in Tcas5.2 is also listed.

| ChLG | Tcas5.0 scaffolds | Unplaced scaffolds added in Tcas5.2 | Tcas5.2 super scaffolds |
| --- | --- | --- | --- |
| X | 13 | +2 | 2 |
| 2 | 18 | +1 | 10 |
| 3 | 29 | +4 | 20 |
| 4 | 6 | +2 | 2 |
| 5 | 17 | +1 | 4 |
| 6 | 12 | +6 | 6 |
| 7 | 15 | - | 6 |
| 8 | 14 | +1 | 8 |
| 9 | 21 | - | 9 |
| 10 | 12 | +2 | 10 |
| Total | 157 | 19 | 77 |

**Figure 8.**
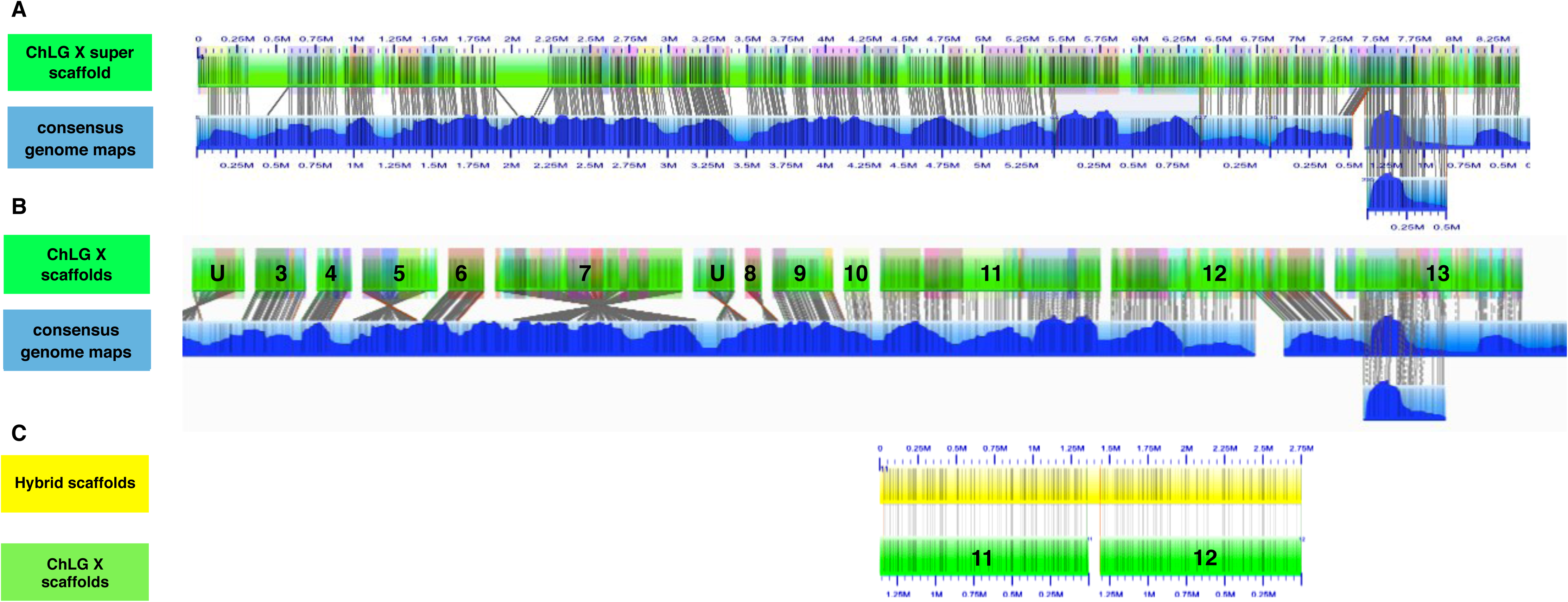
ChLGX before and after super scaffolding with Stitch and Hybrid Map creation by Hybrid Scaffold. (A) Alignment of Tcas5.2 *in silico* maps to consensus genome maps for ChLGX. (B) Alignment of Tcas5.0 *in silico* maps to consensus genome maps for ChLGX. (c) Alignment of Hybrid genome maps to consensus genome maps for ChLGX. Consensus genome maps are blue with molecule map coverage shown in dark blue). The *in silico* maps are green with contigs overlaid as translucent colored squares. Only *in silico* maps that align are displayed in IrysView. The single hybrid scaffold created for ChLGX is shown (yellow). Only new ”hybridized” maps are shown in IrysView. Labels and alignments of labels are indicated with grey lines.

The Tcas5.2 Whole Genome Shotgun project has been deposited at DDBJ/EMBL/GenBank under the accession AAJJ00000000. The version described in this paper is version [DDBJ:AAJJ02000000,EMBL:AAJJ02000000,GenBank:AAJJ02000000]. Two scaffolds were removed from the genome assembly because they were identified as contaminants after they blasted to the *Bos frontalis* genome.

### Assembly: Putative Haplotypes

Evidence of putative haplotypes was found during visual inspection of alignments. Although overall alignment redundancy was rare for the *Strict*-*T* assembly; when observed, it usually consisted of only two consensus genome maps aligning to the same *in silico* map (Figure 9). This redundancy might indicate collapsed repeats in the sequence-based assembly. On the other hand it could also indicate segmental duplications, assembly of alternative haplotypes, or mis-assembly producing redundant consensus genome maps. In Figure 9A, two consensus genome maps aligned to the same *in silico* map. One consensus genome map aligned across most of the *in silico* map while only a small region of the other consensus genome maps aligned to the the *in silico* map (Figure 9A). Molecule map coverage decreased for each consensus genome map in the region where only one of them aligned to the *in silico* map (Figure 9A-C). Taken together, the region of lower coverage and the number of consensus genome maps aligning (two) may indicate the assembly of two haplotypes.

**Figure 9.**
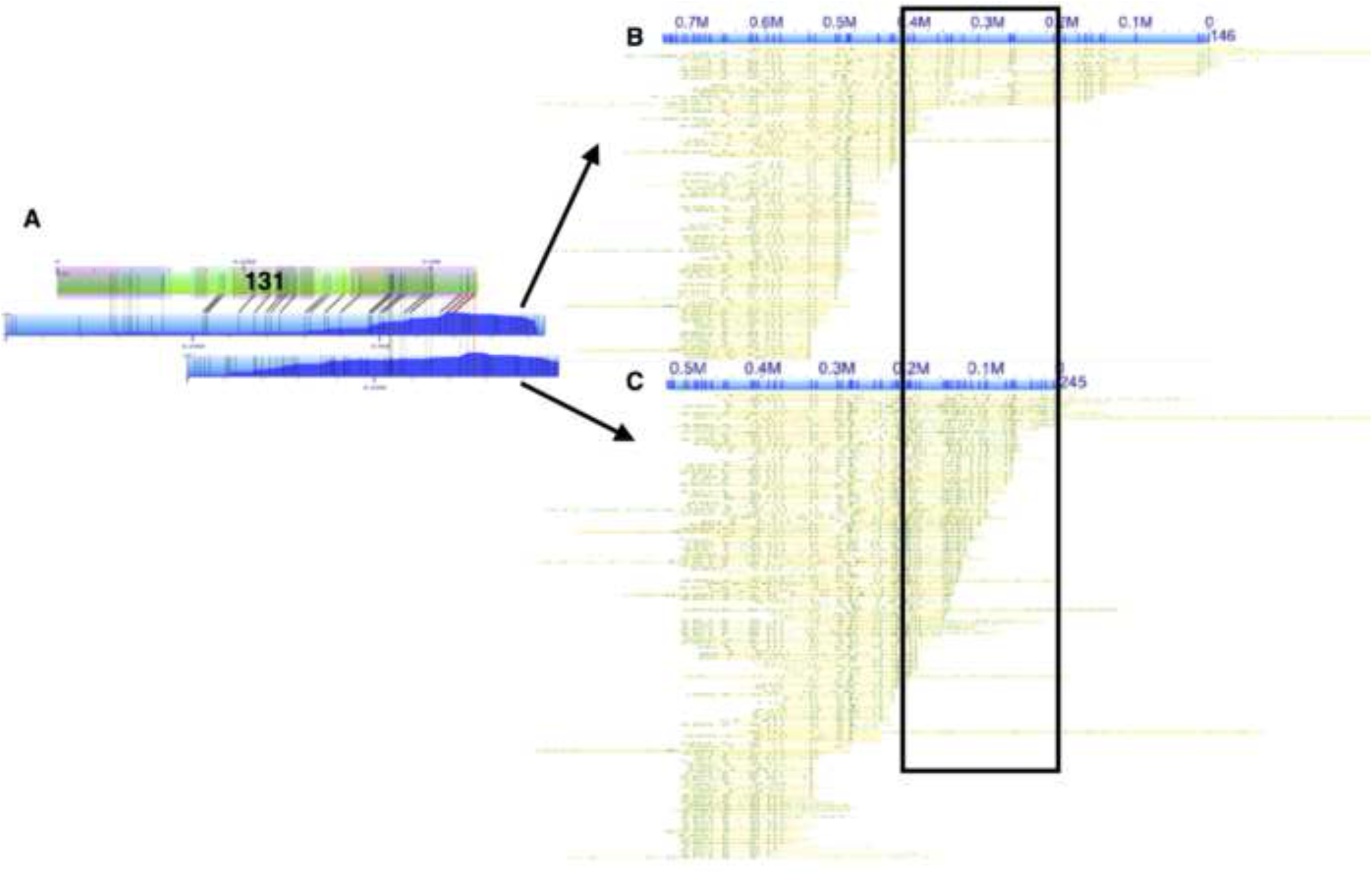
Putative haplotypes assembled as consensus genome maps. (A) Two consensus genome maps (blue with molecule map coverage shown in dark blue) align to the *in silico* map of scaffold 131 (green with contigs overlaid as translucent colored squares). (B and C) Both consensus genome maps are shown (blue) with single molecule pileups (yellow). Both consensus genome maps have similar label patterns except within the lower coverage region indicated with a black square.

### Stitch: Comparison to Other Software

We also ran Hybrid Scaffold (BioNano version 3076) to improve the *T. castaneum* sequence assembly using the *T. castaneum* consensus genome maps and the *in silico* maps from Tcas5.0. Although Hybrid Scaffold improved the scaffold N50 of the sequence-based assembly from 1.16 Mb to 1.83 Mb, the increase in N50 was not as great as the increase after running Stitch (N50 = 3.85 Mb) (Table 2). The total number of scaffolds in the sequence assembly decreased by 30 after running Hybrid Scaffold; in comparison, Stitch reduced the number of scaffolds by 92. Stitch increased the length of the assembly by 4.98 Mb while Hybrid Scaffold increased assembly length by 14.80 Mb. The additional increase in length from Hybrid Scaffold is likely due to extension of sequence-based scaffolds with end gaps introduced from genome map contigs that overlap the start or end of a sequence-based scaffold.

Overall, we found Hybrid Scaffold to be more conservative than Stitch. For example in Figure 8B, the alignment of 13 *in silico* maps to three consensus genome maps was input into both Stitch and Hybrid Scaffold. In this alignment, the order of 11 of these *in silico* maps agreed with the order suggested by the genetic map (after reorienting three *in silico* maps). The *in silico* maps 12 and 13 aligned with a negative gap length between them, suggesting they may be mis-assembled or that local assembly may collapse the assembly in this region. There are several possible approaches when considering this kind of conflicting evidence. The approach of Stitch was to record that we have confirmed the relative position of these scaffolds in the larger context of the genome by creating a new super scaffold containing 100 bp spacer gaps to indicate that exact overlap or gap length is unclear (Figure 8A). Stitch also reports the negative gap length to indicate the need for further sequence level evaluation at a later date. Alternatively, Hybrid Scaffold only automates genome improvements that are unambiguously supported by all lines of evidence (sequence-based and genome map-based) and leaves any ambiguous decisions to a human curator. Therefore, Hybrid Scaffold would only report the conflicting alignment. This is why scaffold 13, for example, is not included in the hybrid map produced by Hybrid Scaffold (Figure 8C).

For highly refined genome assemblies, this lack of tolerance for noisy alignments has not hindered improvement of genome projects. However, for less refined genome assemblies this may be too stringent. The Hybrid Scaffold software, for example, was developed to scaffold the human genome and has been found to work well for this application. Genome projects at earlier stages would benefit from staged release of updates (e.g., immediate release of general improvement in scaffold order and orientation followed by further refinement at the sequence level). For projects such as *T. castaneum*, a more aggressive algorithm such as Stitch may be preferred in order to release the bulk of the new information about the higher order arrangement of the genome to the community of *T. castaneum* researchers quickly. Further refinement at a sequence level can be released in subsequent genome updates as they are completed.

### Stitch: Assembly and Super Scaffolding With Multiple Genera

We examined experiments from 16 different genera to determine if the results seen for the *Tribolium castaneum* genome are typical for other genomes as well. The *T. castaneum* genome map N50 was found to be in the high end of the probability density distribution (Additional file 6; Section 1). The same is true for the Tcas5.0 draft sequence assembly N50 and percent of N50 improvement after super scaffolding compared to the other 17 of 19 total projects that had draft sequence genomes (Additional file 6; Section 1). However, in no case was the *T. castaneum* value the highest value recorded, suggesting that a wide range of output quality is possible including values better and worse than the output for *T. castaneum*.

We checked for evidence of correlations between a range of genomic metrics and map assembly, alignment or FASTA super scaffolding results. Because many of the genomic metrics had very broad ranges with variance that increased often for higher values the genomic metrics were log transformed to compress the upper tails and stretch the lower tails of the distributions.

Overall we found little correlation between either sequence FASTA N50, molecule map coverage or molecule map label density and final genome map N50. We did, however, find correlations between finished map and sequence assembly metrics and alignment and super scaffolding quality. There is a positive correlation between high value sequence assembly metrics and *in silico* map-to-genome map alignment metrics (Additional file 6; Sections 3-5) as well as post super scaffolding N50 improvement (Additional file 6; Sections 3,5). There is also a positive correlation between high value genome map assembly metrics and post super scaffolding N50 improvement (Additional file 6; Sections 3,5). However, no direct correlation was found between sequence assembly N50 and genome map N50 (Additional file 6; Sections 4-5). Taken together the analysis suggests different factors may determine sequence assembly and genome map assembly quality. Although sequence assembly N50 may not be useful to predict genome map N50, if both independent assemblies have high N50’s than more of the map lengths may align and super scaffolding may be more productive.

The low degree of correlation found between genome map N50 and sequence N50 may stem from steps unique to the molecule map imaging process. It might be expected that a genome with sequence that assembles well may have qualities that would also favor molecule map assembly (e.g. low repeat content, low ploidy, inbreed lines, etc.). However molecule map assembly is also influenced by unique factors like frequency of fragile sites (two labels occurring on opposite strands in close proximity), labeling efficiency and ability to extract high molecular weight DNA all of which vary for different organisms.

Principal component analysis (PCA) suggests a negative correlation between labels per 100 kb and molecule coverage (Additional file 6: Sections 2-3). The correlation between labels per 100 kb and molecule coverage was weakly significant in individual regression (Additional file 6; Sections 4-5). Labels per 100 kb are monitored as molecules are being imaged. Lower than expected label density can occasionally lead to further labeling reactions or other adjustments to data collection and therefore greater depth of coverage.

Overall, comparison of the results for the *T. castaneum* genome and 19 additional genome projects suggest that results may vary widely from project to project. Many factors may contribute to this effect including the quality of the sequence assembly, degree of divergence between the organism or organisms used to extract DNA, success of extraction and labeling of high molecular weight DNA, genome size and genome complexity. In fact, the tendency for assemblies from the same genera or species to cluster together on the PCA plots suggests that organism-specific qualities may influence assembly, alignment or super scaffolding results (Additional file 6: Sections 2-3). Although analysis of more projects is needed to determine if these similarities are meaningful predictors of output quality.

## Conclusions

We introduced new tools to facilitate single molecule map assembly optimization and genome finishing steps using the resultant consensus genome maps. These tools were validated using the medium-sized (200 Mb) *T. castaneum* beetle genome. The Tcas3.0 genome was assembled using the gold-standard Sanger assembly strategy [15]. The Tcas5.0 assembly benefitted from the use of LongDistance Illumina Jump libraries to anchor additional scaffolds and fill gaps. Despite this, we were able to more than triple the scaffold N50 by leveraging the optimal consensus genome maps and Stitch. We demonstrated that the AssembleIrysCluster method of optimization and Stitch can be used together to improve the contiguity of a draft genome.

As the variety of genome assembly projects increases, we are discovering that tools appropriate for all projects (e.g. genomes of varying size and complexity, assemblies of varying quality, various taxonomic groups, etc.) do not exist. Indeed, the results of Assemblation2 indicate that no one suite of datatypes or assembly workflow may be sufficient to best assemble even the subcategory of vertebrate genomes [3]. Here we described two software tools and many shorter scripts to summarize and work with these new data formats. However, we anticipate the development of a variety of bioinformatics tools for extremely long, single molecule map data as more applications for these maps are explored.

Some draft assemblies may currently be too fragmented to align to genome maps assembled from single molecule maps. However as NGS genome assemblies improve from longer read advancements existing genome maps may become useful for scaffolding new or updated sequence assemblies.

Regions where consensus genome maps disagree with sequence assemblies (e.g. negative gap lengths or partial alignments) are flagged by Stitch for investigation at a sequence level. Bioinformatics tools that could automate assembly editing based on such discrepancies are needed to fully support genome improvement with consensus genome maps.

## Availability and requirements

### Pipelines and tutorials

**Project name:** Sewing machine pipeline

**Project home page:** The Sewing machine script and tutorial are available at https://github.com/i5K-KINBRE-script-share/Irys-scaffolding/blob/master/KSU_bioinfo_lab/stitch

**Operating system(s):** Linux (tested on CentOS 7, Gentoo and Ubuntu).

**Programming language:** Perl, Rscript, Bash

**License:** Pipeline script and tutorial are available free of charge to academic and non-profit institutions.

**Any restrictions to use by non-academics:** Please contact authors for commercial use.

**Dependencies:** Sewing machine requires BioPerl and BNGCompare. RefAligner is also required between iterations and can be provided by request by Bionano Genomics http://www.bionanogenomics.com/.

**Project name:** “ Raw data-to-finished assembly and assembly analysis” pipeline

**Project home page:** The pipeline script and tutorial are available at https://github.com/i5K-KINBRE-script-

**Operating system(s):** Xeon Phi server with 1488 threads (6x60x4 Xeon Phi co-processor threads + 24x2 Xeon host threads) and 256GB of host RAM + 6 x 8GB Xeon Phi Ram, and Linux CentOS 7.

**Programming language:** Perl, Rscript, Bash

**License:** Pipeline script and tutorial are available free of charge to academic and non-profit institutions.

**Any restrictions to use by non-academics:** Please contact authors for commercial use.

**Dependencies:** AssembleIrysXeonPhi.pl and AssembleIrysCluster.pl requires DRMAA job submission libraries. RefAligner and Assembler are also required and can be provided by request by Bionano Genomics http://www.bionanogenomics.com/.

**Project name:** “Raw data-to-finished de novo assembly and assembly analysis” pipeline Project home page: The pipeline script and tutorial are available at https://github.com/i5K-KINBRE-script-share/Irys-scaffolding/blob/master/KSU_bioinfo_lab/assemb

**Operating system(s):** Xeon Phi server with 1488 threads (6x60x4 Xeon Phi co-processor threads + 24x2 Xeon host threads) and 256GB of host RAM + 6 x 8GB Xeon Phi Ram, and Linux CentOS 7.

**Programming language:** Perl, Rscript, Bash

**License:** Pipeline script and tutorial are available free of charge to academic and non-profit institutions.

**Any restrictions to use by non-academics:** Please contact authors for commercial use.

**Dependencies:** AssembleIrysXeonPhi.pl and AssembleIrysCluster.pl requires DRMAA job submission libraries. RefAligner and Assembler are also required and can be provided by request by Bionano Genomics http://www.bionanogenomics.com/.

### Assembly scripts

**Project name:** AssembleIrysXeonPhi.pl / AssembleIrysCluster.pl

**Project home page:** AssembleIrysXeonPhi scripts are available at https://github.com/i5K-KINBRE-script-The currently unsupported AssembleIrysCluster scripts are available on Github at

https://github.com/i5K-KINBRE-script-share/Irys-scaffolding/tree/master/KSU_bioinfo_lab/assemb

**Operating system(s):** Xeon Phi server with 1488 threads (6x60x4 Xeon Phi co-processor threads + 24x2 Xeon host threads) and 256GB of host RAM + 6 x 8GB Xeon Phi Ram, and Linux CentOS 7 and SGE Linux (tested on a Gentoo) cluster respectively

**Programming language:** Perl, Rscript, Bash

**License:** AssembleIrysXeonPhi and AssembleIrysCluster.pl is available free of charge to academic and non-profit institutions.

**Any restrictions to use by non-academics:** Please contact authors for commercial use.

**Dependencies:** AssembleIrysXeonPhi.pl and AssembleIrysCluster.pl requires DRMAA job submission libraries. RefAligner and Assembler are also required and can be provided by request by Bionano Genomics http://www.bionanogenomics.com/.

### Super scaffolding scripts

**Project name:** stitch.pl

**Project home page:** Stitch scripts are available on Github at https://github.com/i5K-KINBRE-script-shar

**Operating system(s):** MAC and LINUX (tested on Gentoo and Ubuntu)

**Programming language:** Perl, Rscript, Bash

**License:** stitch.pl is available free of charge to academic and non-profit institutions.

**Any restrictions to use by non-academics:** Please contact authors for commercial use.

**Dependencies:** stitch.pl requires BioPerl. RefAligner and Assembler are also required between iterations and can be provided by request by Bionano Genomics http://www.bionanogenomics.com/.

### Map summary scripts

**Project name:** BNGCompare.pl, bnx stats.pl, cmap stats.pl and xmap stats.pl

**Project home page:** all scripts are available on Github at https://github.com/i5K-KINBRE-script-share/I and https://github.com/i5K-KINBRE-script-share/BNGCompare

**Operating system(s):** MAC and LINUX (tested on Gentoo and Ubuntu)

**Programming language:** Perl, Rscript, Bash

**License:** bnx stats.pl, cmap stats.pl and xmap stats.pl are available free of charge to academic and non-profit institutions.

**Any restrictions to use by non-academics:** Please contact authors for commercial use.

**Dependencies:** bnx stats.pl, cmap stats.pl and xmap stats.pl have no dependencies.

### Abbreviations

NGS - next-generation sequencing

BAC - bacterial artificial chromosome

QC - quality control

PAT - percent aligned threshold

ChLG - chromosome linkage groups

PCA - principal component analysis

### Competing interests

The JMS, MCC, NH, NL, and SJB declare that they have no competing interests. ETL, PS and TA are employees at BioNano Genomics and hold stock options.

### Author’s contributions

MCC isolated the high molecular weight DNA and generated the image files on the Irys. ETL and JMS developed the assembly workflow. JMS wrote most of the code in the IrysScaffolding Github Repo (Stitch, AssembleIrysXeonPhi, AssembleIrysCluster, etc.). NH assisted with initial code review of analyze irys output (precursor to Stitch) and prepared Tcas5.0. JMS and NL manually edited Tcas5.1. JMS performed the data analyses. TA contributed to sections discussing BioNano RefAligner and Assembler. PS contributed to interpretation of results. JMS and SJB did most of the writing with contributions from all authors. All authors read and approved the final manuscript.

## Acknowledgements

This project was supported by an Institutional Development Award (IDeA) from the National Institute of General Medical Sciences of the National Institutes of Health under grant number P20 GM103418. The content is solely the responsibility of the authors and does not necessarily represent the official views of the National Institute of General Medical Sciences or the National Institutes of Health.

Data for Additional file 1 and Additional file 6 was kindly made available by P.A. Larsen, J. Rogers, A.D. Yoder and the Duke Lemur Center.

The Tribolium castaneum genome project is part of the i5k Genome Sequencing Initiative for Insects and Other Arthropods.

Matthias Weissensteiner & Jochen Wolf, Uppsala University. Stephen Schaeffer from The Pennsylvania State University and Stephen Richards from the Baylor College of Medicine Human Genome Sequencing Center for the use of the D. pseudoobscura data. Mike Kanost from Kansas State University. Jeff Maughan from Brigham Young University for the use of the Amaranth data. The Udall Lab from Brigham Young University and Cotton Inc. for the use of the cotton data. Grant (NSF 1237993) for use of the Medicago data. Christopher Cunningham, University of Georgia for the use of Nicrophorus data. Catherine Peichel from the Fred Hutchinson Cancer Research Center and Michael White from the University of Georgia for the Gasterosteus data. Mirkό Palla, Ph.D., Wyss Institute Postdoctoral Fellow, Church; Laboratory - Department of Genetics, Harvard Medical School and George Church, Ph.D., Wyss Institute Core Faculty Member, Robert Winthrop Professor of Genetics at Harvard Medical School, Professor of Health Sciences and Technology at Harvard and MIT, and Senior Associate Member at the Broad Institute of Harvard and MIT for the Escherichia coli data.

## Additional Files

Additional file 1 — Single molecule map stretch per scan in recent flowcells.

Bases per pixel (bpp) is plotted for scans 1..*n* for each flowcell of mouse lemur molecules (purple). The first scan of each flowcell is indicated with a grey dashed line. The pre-adjusted molecule map stretch was determined by aligning molecule maps to the *in silico* maps. Data made available by P.A. Larsen, J. Rogers, A.D. Yoder and the Duke Lemur Center.

Additional file 2 — Cumulative length and number of single molecule maps per BNX file for *T. castaneum* data generated over time

Detailed metrics for molecule maps per BNX file (cumulative length and number of maps). Columns include cumulative length of molecule maps > 150 kb, number of molecule maps > 150 kb and date that BNX file was generated.

Additional file 3 — Single molecule map metrics and histograms from *T. castaneum* DNA

Detailed metrics for molecule maps including map N50, cumulative length and number of maps. Figures show histograms of per molecule map quality metrics including length, molecule map SNR and intensity, label count, label SNR and label intensity. Molecule maps are filter for minimum molecule lengths of 100, 150 or 180 kb.

Additional file 4 — Assembly of *T. castaneum* consensus genome maps with range of parameters

Detailed assembly metrics for assembled consensus genome maps using strict, default and relaxed ”-T” parameter, p-value threshold are named Relaxed-T, Default-T and Strict-T respectively. The best ”-T” parameter was used for two additional assemblies with either relaxed minimum molecule map length (relaxed-minlen) of 100 kb, rather than the 150 kb default, or a strict minimum molecule map length (strict-minlen) of 180 kb.

Additional file 5 — ChLGs before and after super scaffolding

Alignments of Tcas5.0 and Tcas5.2 *in silico* maps to consensus genome maps for all ChLGs. Consensus genome maps (blue with molecule coverage shown in dark blue) aligned to the *in silico* maps (green with contigs overlaid as translucent colored squares). Alignment to both Tcas5.2 super scaffolds (top alignment) and Tcas5.0 scaffolds (bottom alignment) are shown.

Additional file 6 — Assembly and super scaffolding with multiple genera.

We examined experiments from 16 different genera to determine if the results seen for the Tribolium castaneum genome are typical for other genomes as well.

